# Inflammation induced by influenza virus impairs innate control of human pneumococcal carriage

**DOI:** 10.1101/347161

**Authors:** Simon P. Jochems, Fernando Marcon, Beatriz F. Carniel, Mark Holloway, Elena Mitsi, Emma Smith, Jenna F. Gritzfeld, Carla Solórzano, Jesús Reiné, Sherin Pojar, Elissavet Nikolaou, Esther L. German, Angie Hyder-Wright, Helen Hill, Caz Hales, Wouter A.A de Steenhuijsen Piters, Debby Bogaert, Hugh Adler, Seher Zaidi, Victoria Connor, Jamie Rylance, Helder I. Nakaya, Daniela M. Ferreira

## Abstract

Secondary bacterial pneumonia following influenza infection is a significant cause of mortality worldwide. Upper respiratory tract pneumococcal carriage is important as both determinants of disease and population transmission. The immunological mechanisms that contain pneumococcal carriage are well-studied in mice but remain unclear in humans. Loss of this control of carriage following influenza infection is associated with secondary bacterial pneumonia during seasonal and pandemic outbreaks. We used a human type 6B pneumococcal challenge model to show that carriage acquisition induces early degranulation of resident neutrophils and recruitment of monocytes to the nose. Monocyte function associated with clearance of pneumococcal carriage. Prior nasal infection with live attenuated influenza virus induced inflammation, impaired innate function and altered genome-wide nasal gene responses to pneumococcal carriage. Levels of the cytokine IP-10 promoted by viral infection at the time of pneumococcal encounter was positively associated with bacterial density. These findings provide novel insights in nasal immunity to pneumococcus and viral-bacterial interactions during co-infection.

## Main

Pneumonia is a major global health problem; it kills more children under 5 years of age than any other disease ^1^. The burden of disease is aggravated by old age, chronic lung disease, immunosupression and viral co-infection. Secondary pneumonia following pandemic and seasonal influenza virus infection is a significant cause of mortality worldwide ^2^.

Nasopharyngeal carriage of *Streptococcus pneumoniae* (Spn, pneumococcus) is common with 40-95% of infants and 10-25% of adults colonised at any time ^3^. Carriage is important as the pre-requisite of infection ^4^, the primary reservoir for transmission ^5^ and the predominant source of immunising exposure and immunological boosting in both children and adults ^6,7^.

Immune dysregulation caused by respiratory virus infection such as influenza leads to increased carriage density ^8^. Increased carriage density has been associated with pneumonia incidence and severity, as well as with within-household Spn transmission ^5,9–11^. The mechanisms and markers associated with this pathogen synergy have been difficult to study in human subjects due to the rapid nature of the disease.

One safe way to simulate influenza infection in the nose is using Live Attenuated Influenza Vaccine (LAIV), consisting of cold-adapted influenza viruses. LAIV has been shown to affect the subsequent susceptibility to Spn and to lead to increased carriage density in murine models of infection and in vaccinated children ^12,13^. In a randomised controlled trial, we showed that LAIV administration prior to Spn challenge led to 50% increase in Spn acquisition by molecular methods as well as 10-fold increase in nasopharyngeal bacterial load ^14^.

In murine models of pneumococcal carriage, Th17-dependent recruitment of neutrophils and monocytes to the nasopharynx mediates immunological control and clearance ^15–17^. Influenza virus infection promotes Type I interferons which interfere with recruitment of these phagocytes, while IFN-L is postulated to impair phagocytosis by macrophages through downregulation of the scavenger receptor MARCO ^18–20^. However, the precise immune mechanisms and gene regulators involved in the control and clearance of pneumococcal carriage in humans have not been revealed ^21^. Moreover, how these mechanisms are altered during human influenza virus infection remains largely unknown.

In recent years, systems biology approaches have allowed for the identification of immune mechanisms associated with protection from infectious diseases and with robust immune responses during vaccination ^22–28^. Here, we applied systems biology to nasal samples collected in the setting of human challenge with LAIV and Spn, to emulate nasal effects of influenza infection on Spn carriage. We identified for the first time in humans the key cellular mechanisms that control newly acquired pneumococcal carriage, and how they are disrupted following nasal influenza infection.

## Results

### LAIV-induced inflammation leads to increased pneumococcal carriage density and acquisition

In a double-blinded controlled randomized clinical trial, we administered LAIV (n=55) three days prior to Spn inoculation (day 0). To verify the requisite topical application for an effect on pneumococcal carriage, we administered tetravalent inactivated influenza vaccine (TIV) as a control (n=62). LAIV infection led to transiently increased pneumococcal acquisition at day 2 (60.0% vs 40.3% by molecular methods in LAIV vs control groups, respectively) ^14^. LAIV also increased Spn carriage density in the first 14 days following pneumococcal inoculation (Fig S1 and ^14^). We collected a series of nasal micro-biopsies and nasal lining fluid throughout the study to assess ongoing cellular and cytokine responses. Participants were grouped in those who did not become colonized following Spn challenge (carriage-) and those who did (carriage+), as determined by classical microbiology (Fig 1a). To investigate whether LAIV-induced immune responses were associated with a predisposition to pneumococcal carriage, we measured levels of 30 cytokines in nasal lining fluid (Fig 1b). At day 0, directly prior to Spn inoculation, LAIV significantly increased levels of twenty cytokines after multiple testing correction, including IP-10, TNF-α, IL-10, IFN-◻ and IL-15 (Fig 1b and Table S1). In contrast, the control group did not show any significant increase in cytokine response at day 0. Following Spn inoculation, Spn carriage in the absence of LAIV was associated with increased levels of EGF at day 2 (p = 0.023) and decreased levels of IL-1RA at day 9 post Spn inoculation (p = 0.027) compared to baseline, neither of which remained significant after multiple testing correction. No other cytokines, including IL-17A or MCP-1, were significantly altered by carriage alone (Fig 1b).

**Figure 1.**
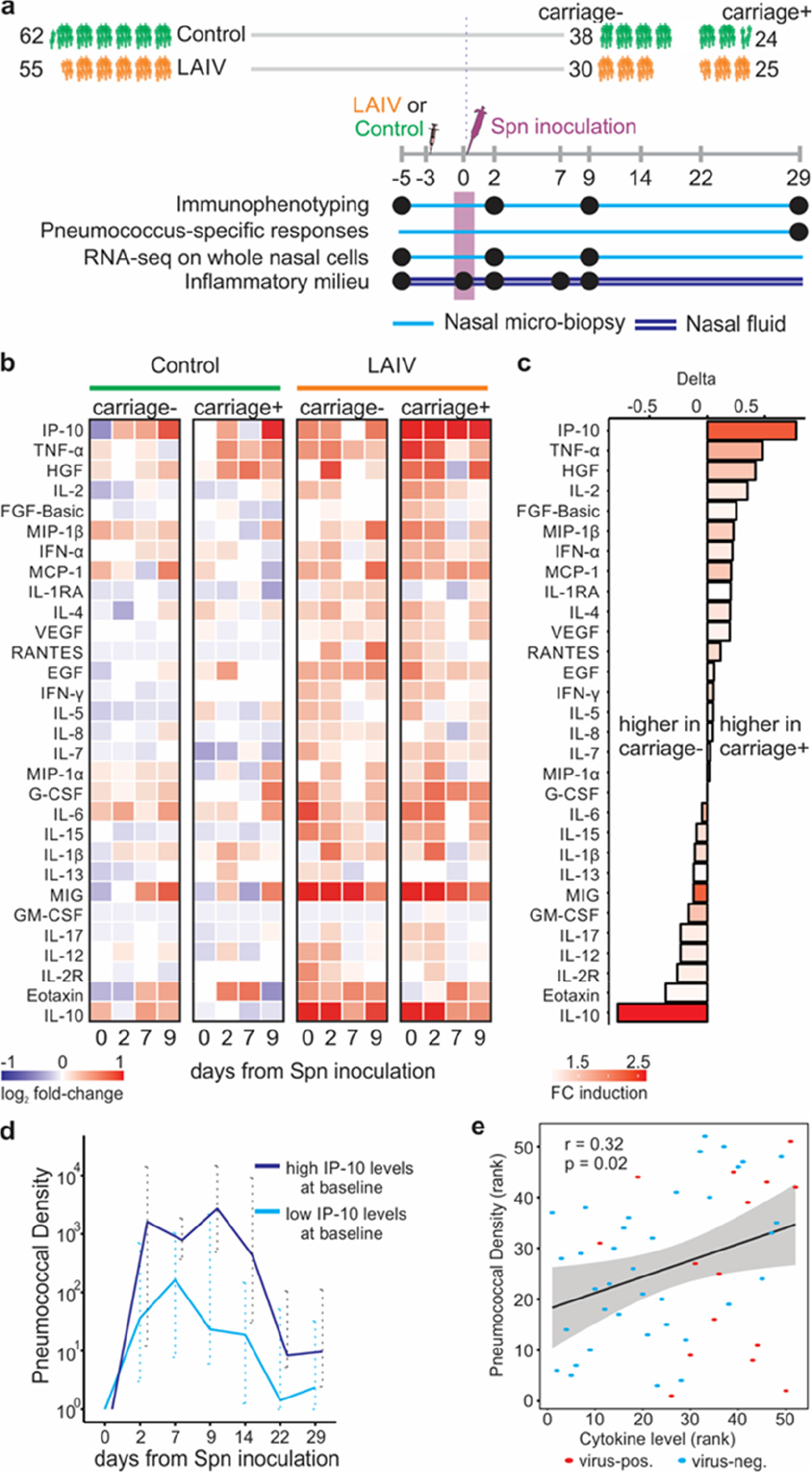
LAIV-pneumococcus co-infection leads to excessive pro-inflammatory responses that associate with increased pneumococcal density and impaired monocyte recruitment. a) Experimental design of the study. LAIV = live attenuated influenza vaccine, Spn = *Streptococcus pneumoniae*. Analysed timepoints are indicated by black circles. b) Heatmap showing for each cytokine the median log2 fold-change compared to baseline for the timepoints 0/2/7/9, n = 19 per group. c) The delta in median log fold change (FC) following LAIV vaccination just prior to inoculation with Spn for subjects becoming carriage+ or carriage-excluding subjects becoming positive by PCR only, who resemble subjects that become carriage+ by culture as well). The colour of each bar represents the median induction in the entire LAIV group. d) Pneumococcal density (median and interquartile range of CFU/mL nasal wash shown) for all carriage+ subjects with high (top quartile) or low (all subjects below top quartile) IP-10 levels at day 0. P = 0.019 by Mann-Whitney of AUC of log-transformed density over time. e) Scatter plot showing correlation of levels of IP-10 at baseline with Spn density for a second validation cohort (n = 52) with an asymptomatic upper respiratory tract virus infection (n=15) or not. Spearman correlation test results are shown.

Even before bacterial inoculation, nasal inflammatory responses to LAIV differed between those who went on to become carrier and those who were protected from carriage (Fig 1c). In particular, IL-10 was increased more in LAIV-vaccinated subjects who did not acquire Spn following inoculation (5.8-fold increase, p = 0.0097) than in those who became carriers following inoculation (2.0-fold increase, p = 0.073). In contrast, IP-10 was increased more in subjects who went on to become carriers (2.4-fold increase, p = 0.0008) than in those who remained carriage-negative (1.5-fold increase, p = 0.051). Moreover, subjects with increased levels of IP-10 before inoculation displayed higher pneumococcal density following Spn inoculation (Fig 1d). This suggests that differences in the response to influenza virus are associated with secondary susceptibility to Spn. To test whether this was specific for LAIV infection, we measured IP-10 in nasal washes from an independent cohort in which a subset of subjects had asymptomatic viral upper respiratory tract infection before Spn inoculation. These comprised rhinovirus (n=12), coronavirus (n=5), respiratory syncytial virus (n=2) and parainfluenzavirus (n=1) ^29^. The predominant virus, rhinovirus, was recently shown to associate with increased pneumococcal acquisition and transmission ^30^. In these virus-infected subjects, levels of IP-10 were increased (Fig S2), and baseline IP-10 levels correlated with increased pneumococcal density also in this second cohort (r = 0.32, p = 0.02, Fig 1e). While the correlation was modest in this validation cohort suggesting that other host and environmental factors are involved, this is the first time a biomarker predicting Spn density has been identified.

### Early neutrophil degranulation in response to carriage is impaired by LAIV infection

In murine models, neutrophil recruitment after onset of carriage contributes to control of the bacteria ^15^. We observed pre-existent high levels of neutrophils in the human nose and pneumococcal carriage did not lead to significant further recruitment of neutrophils (Fig S3a,b). To investigate whether luminal neutrophils were involved in the early control of carriage, we measured myeloperoxidase (MPO) levels, a marker for neutrophil degranulation ^31^, in nasal wash. Levels were increased (2.2-fold, p < 0.05) at 2 days after challenge in control carriage+ but not carriage-individuals (Fig 2a). This neutrophil activation was impaired in the LAIV group, who displayed high carriage density during early carriage and had increased acquisition compared to controls. Together, this suggests that neutrophil degranulation is important for the initial control of carriage. To investigate whether neutrophils were also impaired systemically following LAIV as reported during wild-type influenza infection ^32^, we isolated blood neutrophils before, and at three days after, LAIV administration from a subset of subjects. We confirmed that opsonophagocytic killing of Spn by blood neutrophils was decreased following LAIV (Fig 2b, p < 0.05). This effect could be mimicked by the addition of TNF-α, but not IP-10, to neutrophils from healthy donors *in vitro*, decreasing killing capacity in a dose-dependent manner (Fig 2c,d). Nanostring expression analysis of 594 genes revealed 10 differentially expressed genes in blood neutrophils 3 days post LAIV (Table S2). Among those were *MAP4K2* (3.2-fold increase, p=3.2×10^−5^), which acts on the TNF-α signal transduction pathway ^33^, and *TIGIT* (3.6-fold increase, p=0.008, Fig 2e). *TIGIT* expression levels also negatively correlated with neutrophil killing capacity (r=-0.73, p=0.02, Fig 2f). TIGIT is an “immune checkpoint” protein, which has been described to promote Treg cell function ^34^, but its expression on neutrophils has not been previously appreciated. Incubation of whole blood with recombinant TNF-α increased TIGIT levels on neutrophil surface within 30 minutes in a dose-dependent manner (Fig 2g).

**Figure 2.**
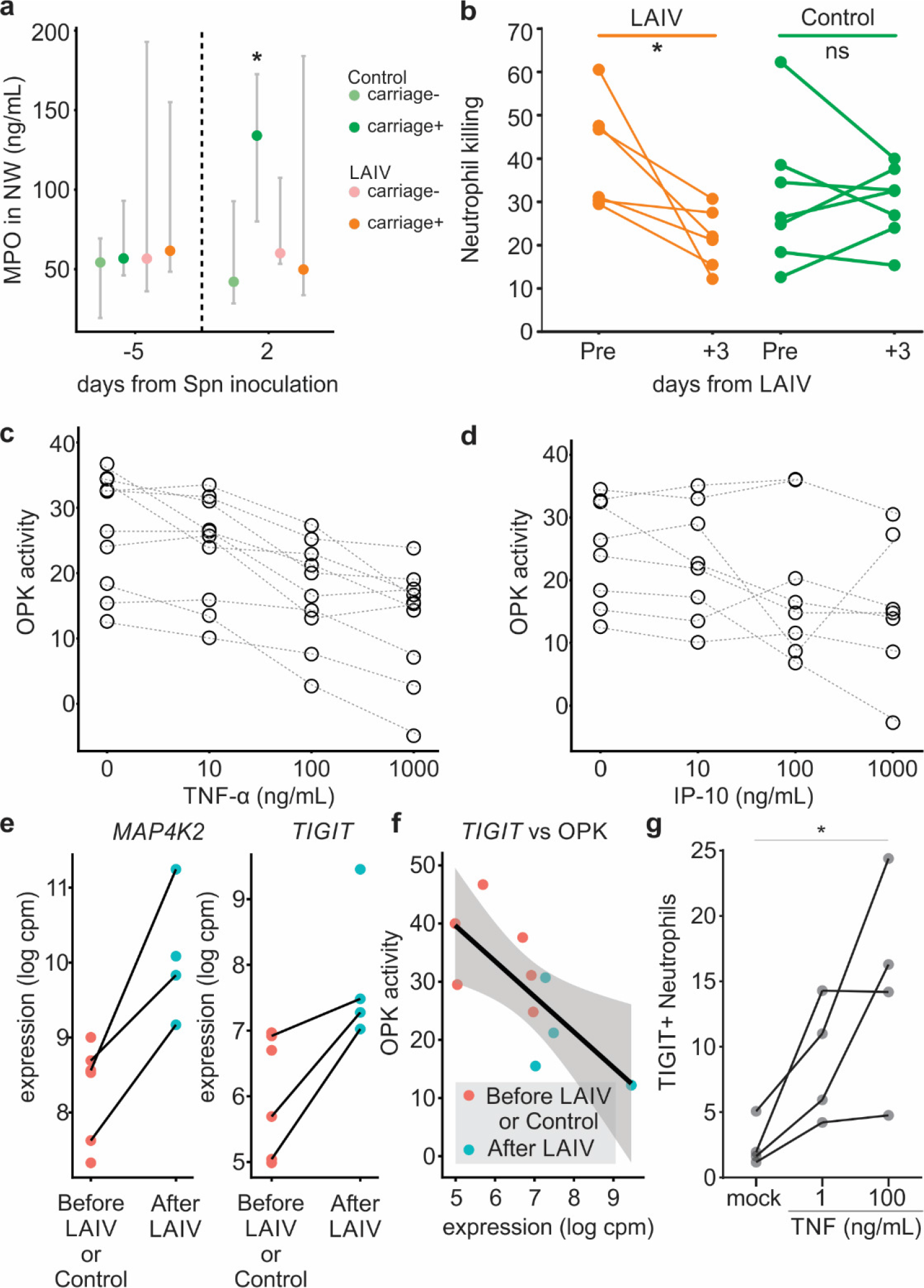
Neutrophil function is impaired following LAIV administration. a) Levels of myeloperoxidase (MPO) in nasal wash of volunteers before or 2 days post Spn inoculation. Median and interquartile range are shown (n=9-10 per group). b) Spn killing capacity of blood neutrophils before and 3 days following LAIV (n=6) or control (TIV or no, n=7) vaccination. Individual subjects are shown and connected by lines. *p < 0.05 by Wilcoxon paired test. c) Effect of exogenous TNF-α and d) IP-10 on the capacity of blood neutrophils of healthy volunteers to kill Spn *in vitro* (opsonophagocytic killing = OPK). Neutrophils from 4-6 subjects were used in 3 independent experiments. Individual samples are depicted and connected by dashed lines. ** p < 0.01, *** p <0.001 by Friedman test followed by Dunn’s multiple comparison. e) Normalized *MAP4K2* and *TIGIT* counts on sorted neutrophils before LAIV or in control arm (red) and following LAIV (blue). Individual samples are shown and paired samples are connected by black lines. ** p < 0.01, *** p < 0.001 f) Correlation between OPK activity and TIGIT counts. Spearman rho and p-value are shown. g) Levels of TIGIT on blood neutrophil surface measured by flow cytometry after a 30 minute incubation without or with 1ng/mL TNF-α or 100ng/mL TNF-α. * p < 0.05, Friedman test followed by Dunn’s multiple comparison.

### Pneumococcal carriage-induced monocytes recruitment to the nose is impaired by LAIV infection

Immunophenotyping revealed a significant recruitment of monocytes to the nose following establishment of carriage (Fig 3a,b and Fig S4). Monocyte levels increased as early as 2 days following Spn inoculation, peaked at 9 days (median 4.8x increase) and remained elevated 29 days post Spn inoculation. In contrast, there was no recruitment of CD3+ T cells to the nose (Fig S4b). LAIV infection prior to pneumococcal carriage impaired the recruitment of monocytes to the nose (Fig 3a,b). Moreover, peak pneumococcal density associated with increased monocyte recruitment in the control group (r=0.51, p=0.02), but not the LAIV group (r=0.08, p=0.70; Fig 3c,d). Indeed, for subjects in the control group with very low carriage densities, which were only detectable by molecular methods, no monocyte recruitment was observed (Fig S4c). This suggests that a minimum Spn load is required for sensing and monocyte recruitment and that LAIV infection interferes with this process. While MCP-1 was not significantly induced following Spn carriage, levels correlated with numbers of monocytes at all timepoints (Fig S5a). Furthermore, stratification of individuals showed that those with increased MCP-1 levels at day 2 post Spn inoculation exhibit increased monocyte recruitment (Fig S5b). Levels of IL-6, IFN-γ and TNF-α also correlated with levels of monocytes at each time point, but stratification of individuals did not reveal a differential recruitment of monocytes (Fig S5a,b). In a second, independent cohort that did not receive any vaccine, monocytes were increased at day 9 following Spn inoculation, which correlated with increased MCP-1 levels in nasal fluid, validating these results (Table S3 and Fig S5c).

**Figure 3.**
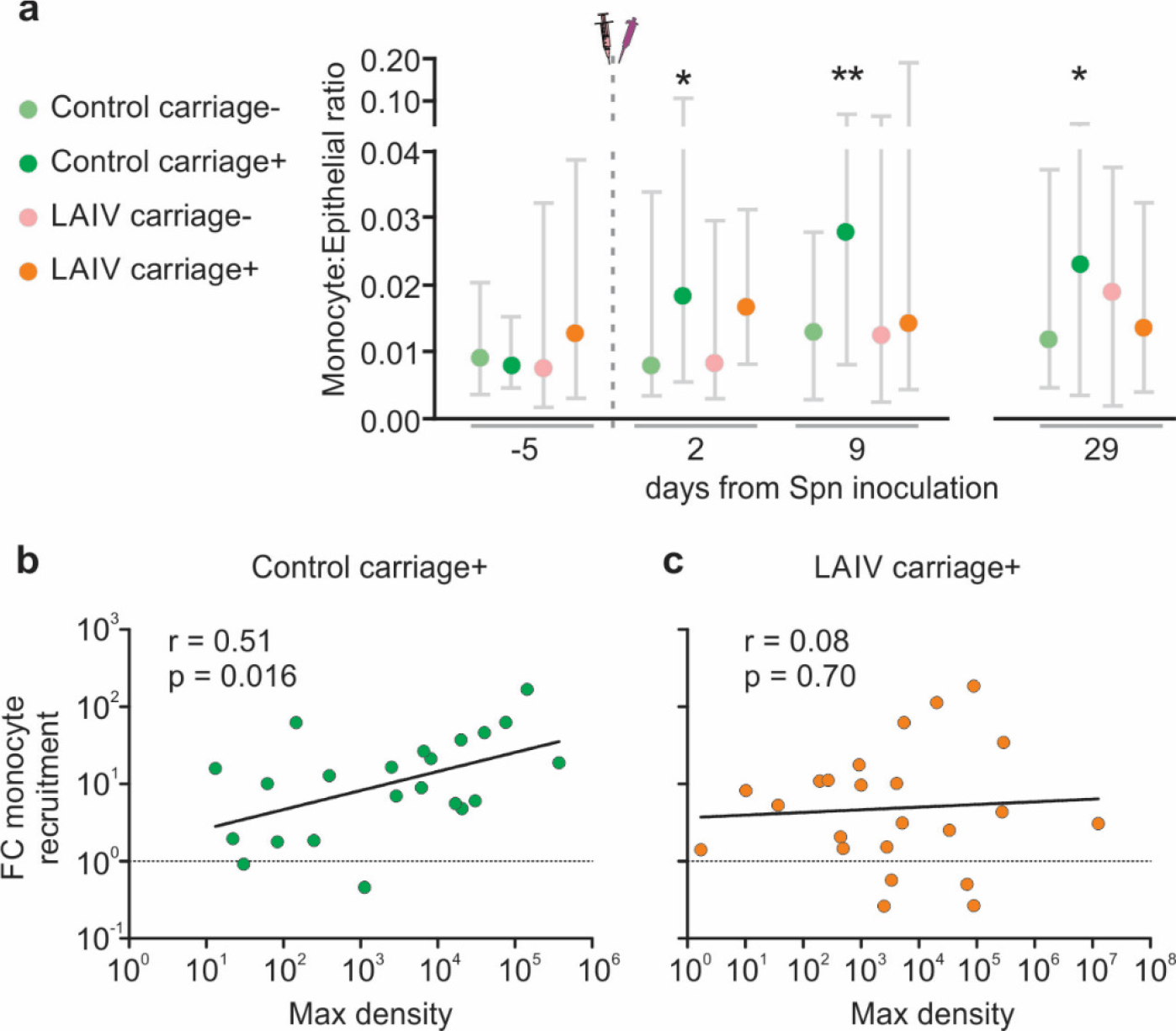
Monocyte recruitment following pneumococcal colonization is impaired during LAIV coinfection. a) Median and interquartile range of nasal monocyte levels normalized to epithelial cell levels are shown per group. b) Representative dot plots showing monocytes for volunteers of the different groups at baseline (day −5) and day of maximum recruitment (day+9). Levels of maximum bacterial density are shown for the c) Control and d) LAIV group and correlated with the maximum monocyte recruitment. Individual subjects are shown and Spearman correlation analysis is shown. *p < 0.05, **p < 0.01 by Wilcoxon paired non-parametric test.

### Nasal responses associated with pneumococcal clearance are impaired by LAIV

To assess anti-pneumococcal responses induced by carriage, we collected nasal cells 29 days post Spn inoculation and stimulated *in vitro* with heat-killed Spn and measured levels of 30 cytokines in supernatant. An increased production (FC > 2 and q < 0.05 to unstimulated control) of TNF-α, MIP-1α, IL-10, IL-6 and GM-CSF upon restimulation was observed in the control carriage+ group (Fig 4a and Fig S6a). In the LAIV carriage+ group, however, this boosting of anti-pneumococcal cytokine responses by re-challenge was absent (Fig 4a and Fig S6a). The production of the above five cytokines correlated with decreased density at day 29 post Spn inoculation, suggesting these responses are involved in Spn clearance (Fig 4b). To test whether monocytes/macrophages were the source of these cytokines we compared the cytokine signature from whole nasal cells with that from alveolar macrophages exposed to Spn *in vitro* (Fig 4c). Relative cytokine production highly correlated (r = 0.66, p < 0.0001) between the two cell populations, suggesting that nasal monocytes/macrophages could be the source of these cytokines. This is supported by the observation that in carriers with low carriage density (only detectable by molecular methods), absence of monocyte recruitment associated with absent Spn-specific responses (Fig S6b)

**Figure 4.**
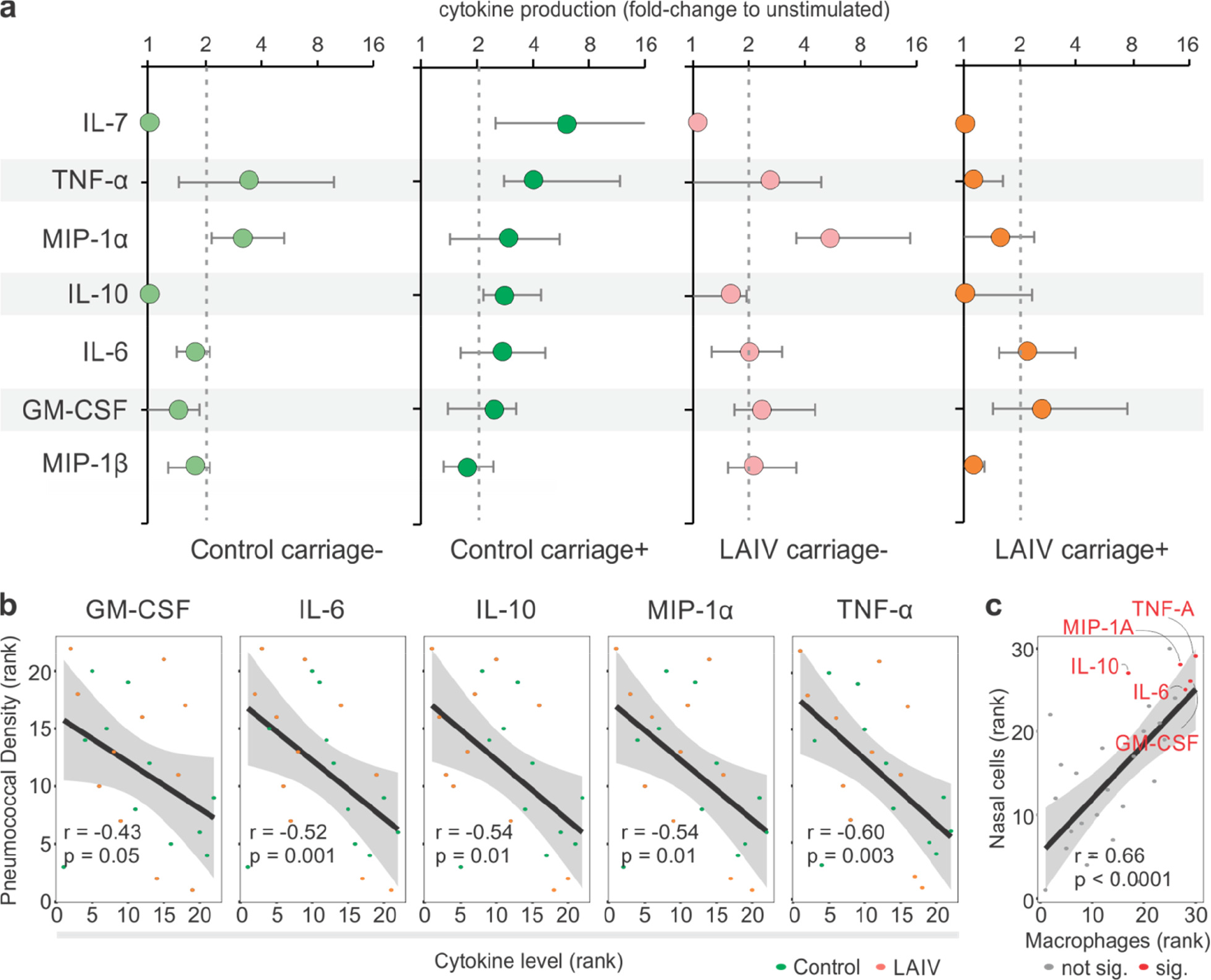
Pneumococcus-specific responses are induced following colonization, which is impaired by LAIV co-infection. a) Whole nasal cells were collected 28 days post-inoculation and stimulated for 18 hours with heat-killed Spn for 48 subjects. Supernatant was collected and levels of 30 cytokines were measured by multiplex ELISA. The median and interquartile range for cytokines induced at least 2-fold in at least one condition are displayed. b) Correlations between cytokine production following Spn stimulation and pneumococcal density are shown. Spearman non-parametric correlation test results are shown per cytokine. c) The cytokine profile from alveolar macrophages (median for 6 volunteers shown) exposed to Spn for 18 hours was compared with that of stimulated whole nasal cells (median of carriage+ group shown).

### LAIV alters nasal gene expression responses to carriage

To identify gene signatures associated with the observed responses to pneumococcal carriage and infection with LAIV, we performed RNA-sequencing on whole nasal cells at days −5, 2 and 9 from Spn inoculation (Fig 5 and Table S4). Carriage without LAIV induced 834 and 176 differentially expressed genes (DEG) at day 2 and day 9, respectively (Fig 5a). These genes were enriched for pathways associated with Gap junction trafficking and regulation (including *GJA1*, *TJP1* and multiple *GJB*) and degradation of the extracellular matrix (including *COL17A1*, *COL12A1*, *LAMA3*, *KLK7*). In the carriage-group, a smaller number of DEG was observed (161 and 248 at day 2 and day 9, respectively).

**Figure 5.**
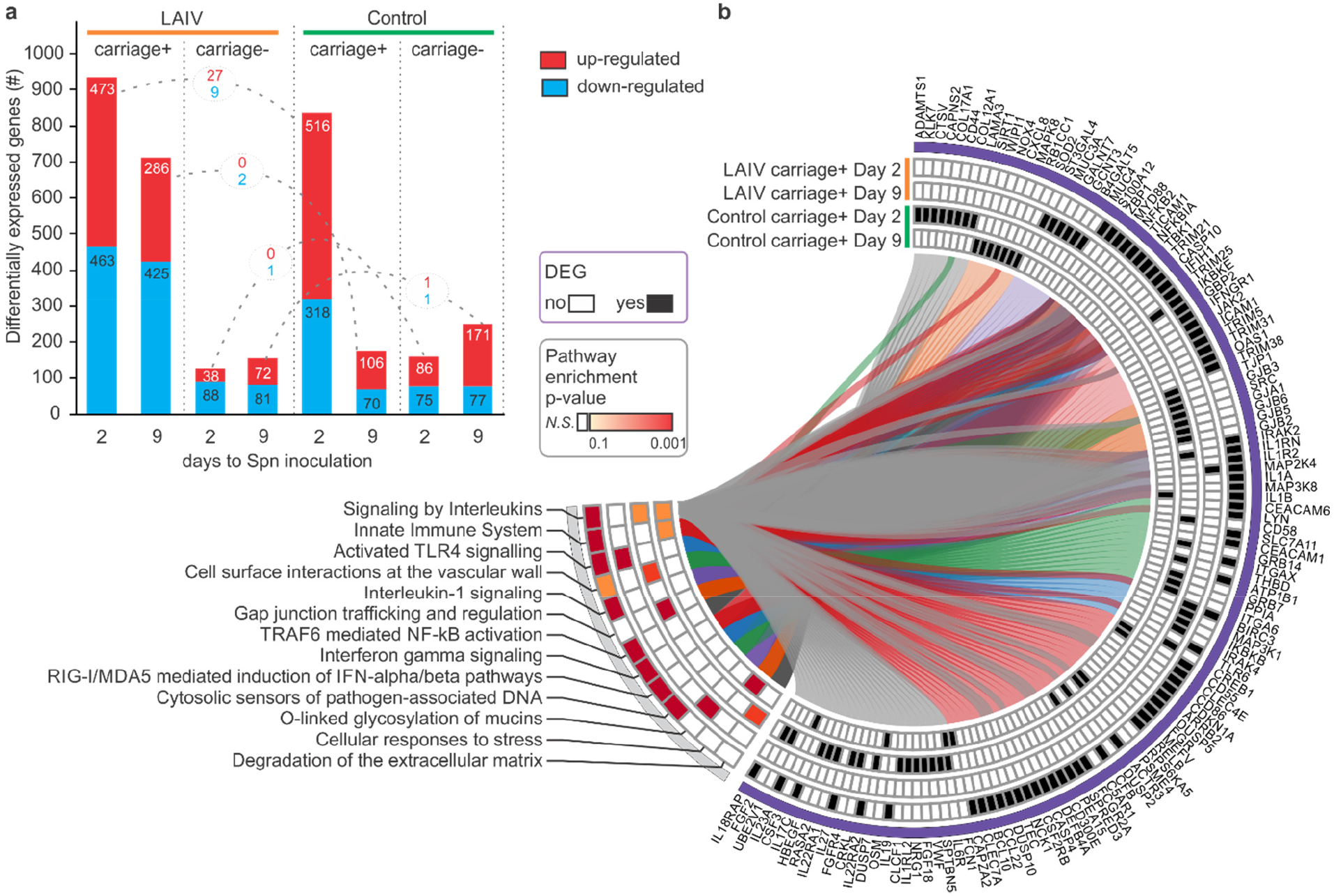
Nasal transcriptomics following LAIV-Spn co-infection (n=35). a) The number of differentially expressed genes (DEGs) between each time point and the baseline for each group are shown. Upregulated and downregulated genes are depicted in red and blue, respectively. Connections between bars show the number of common genes between LAIV and control conditions where colors reflect distinct pathways. b) Circular representation of DEG and Gene Set Enrichment Analysis (GSEA) for LAIV carriage+ and control carriage+ groups at day 2 and day 9 to Spn inoculation. The individual log2-fold changes (baseline-normalized values) values were used as ranks in a single sample GSEA analysis to identify consistently enriched pathways among subjects. Genes and pathways are connected by lines.

In the LAIV carriage+ group, 936 and 711 DEG were observed at day 2 and day 9, respectively. Surprisingly, despite the high levels of inflammatory cytokines observed in the LAIV carriage-group, only a relatively small number of DEG were observed at days 2 and 9 (126 and 153, respectively). DEG of carriage+ subjects receiving LAIV and DEG of carriage+ without LAIV showed very little overlap with only 38 DEG at day 2 and 2 DEG at day 9 in common. Very little overlap was also observed on the pathway level between these groups, indicating LAIV alters the natural responses to pneumococcus (Fig 5b and Table S5). This could reflect transcriptome kinetics, such as observed in altered differentiation and cellular activation, or changes in cell migration to the nasal mucosa.

The LAIV carriage+ group showed an enrichment for genes in TLR3 signalling cascade, RIG-I/MDA5 mediated induction of IFN-alpha/beta pathways and IFN-gamma signalling, which is in agreements with the induction of antiviral responses following LAIV vaccination ^35^. Moreover, TLR4 signalling was also enriched in this group. The pneumococcal protein pneumolysin is sensed by TLR4 ^36^ and it is possible that the increased pneumococcal density following LAIV vaccination led to increased pneumolysin sensing. O-linked glycosylation of mucins, which are used by Spn as a carbohydrate source for growth ^37^, was also enriched in the LAIV carriage+ group (including genes *ST3GAL4*, *GALNT7*, *GCNT3*, *B4GALT5*). *ST3GAL4* is a sialyl transferase and cleavage of sialic acids by the influenza neuraminidase has previously been shown to promote pneumococcal growth ^38^. This finding supports a LAIV-mediated effect on pneumococcal growth through alterations of host factors. Common genes and pathways between the LAIV-vaccinated and control carriers include “Innate immune system” and “Signaling by interleukins” (*IL1B*, *CLEC4E*, *CD55*, *IL1RN*).

### Gene modules associated with recruitment of monocytes

To identify sets of co-expressed genes post LAIV and carriage, we ran our program CEMiTool on the baseline-normalized data of LAIV and control groups, separately ^39^. This modular expression analysis revealed the genes that may act together or are similarly regulated during the immune responses to carriage/infection.

Genes in the control cohort were grouped into four co-expression modules, of which three were significantly enriched for known Reactome pathways (Supp html file 1). Module M1 was enriched in carriage+ at day 9 post Spn inoculation (Fig 6a). Levels of monocytes correlated with the average fold change count in this module, suggesting that these genes reflect the infiltration of monocytes (r = 0.61, p = 0.03, Fig 6b). To further investigate these monocytes, we performed gene set enrichment analysis on the Module M1 genes using list of genes from distinct monocyte subsets (Fig 6c) ^40^. These genes were enriched for classical CD14+CD16-monocytes (p = 0.00002) and not for other monocyte subsets. Moreover, this module was enriched for genes related to “chemokine receptors bind chemokines” and “interferon α/β signalling” (Fig 6d). Type I interferon has been shown to be required for the clearance of pneumococcal carriage in murine models ^41^ and these findings suggest that their activity in monocytes might be critical for this. CEMiTool also integrates co-expression analysis with protein-protein interaction data. Expression of *CXCL6* and its receptor *CXCR2* were identified as hubs in this module M1 (Fig 6e and Supp html file 1). CXCR2 engagement has been shown to induce attachment of monocytes to the endothelial layer, initiating chemotaxis, which suggests this interaction could contribute to monocyte recruitment ^42^. Module M3 was enriched in genes related to “extracellular matrix organization” and “collagen formation” (Fig S7).

**Figure 6.**
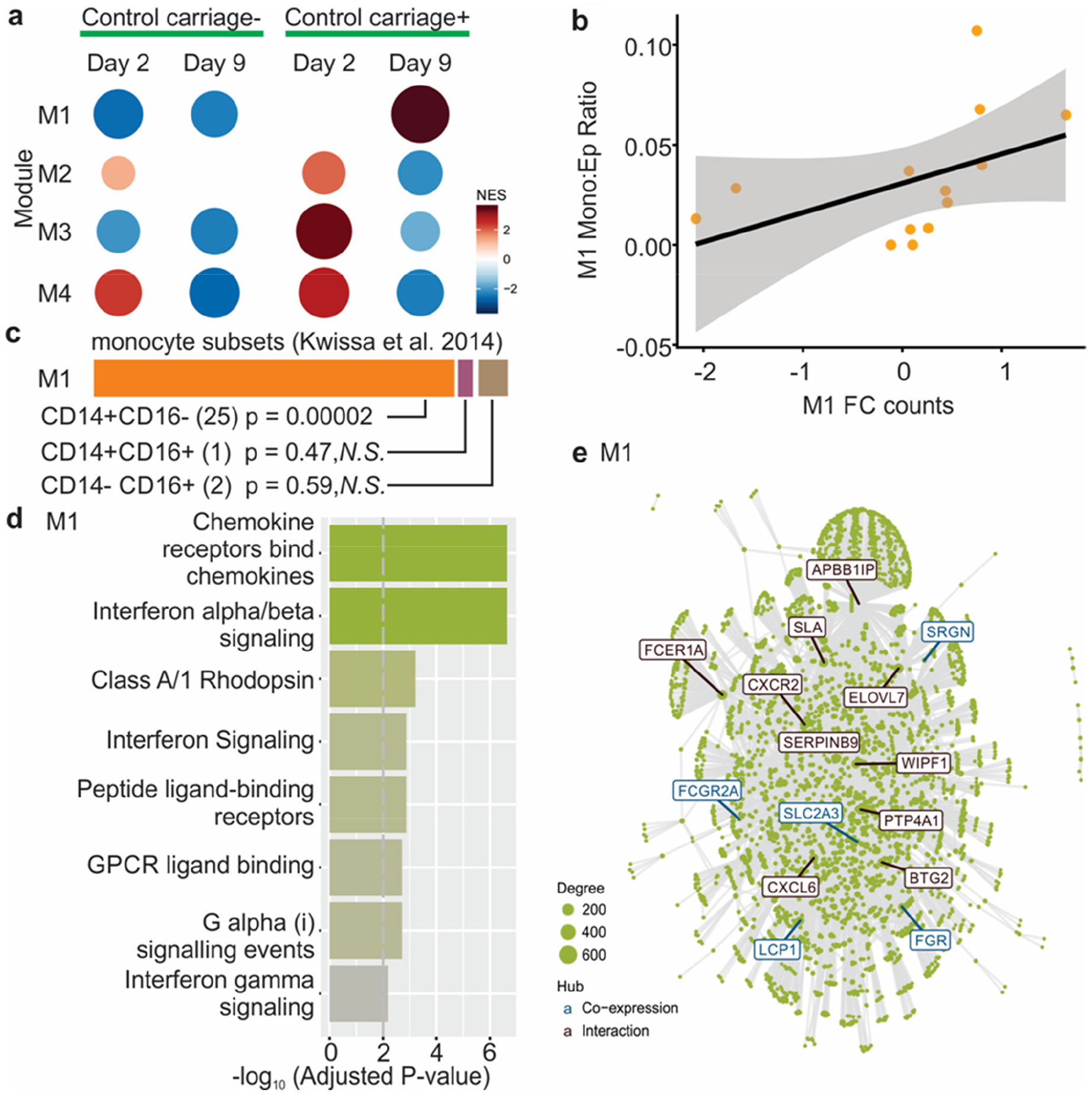
CEMiTool applied to control cohort - Module 1. Raw counts were normalized using log counts per million (CPM) and log2-fold change were calculated for each timepoint against the baseline after which co-expression modules were extracted. a) Gene Set Enrichment Analyses showing the module activity on each timepoint for carriage+ and carriage-groups. b) Correlation with average fold change counts of all M1 genes at day 9 with paired levels of monocytes from the volunteer’s other nostril. Individuals and Spearman correlation analysis are shown. c) The genes of module M1 present with genes highly expressed in CD14+CD16− (578 genes), CD14+CD16+ (108 genes), CD14-CD16+ (162 genes), showing the overlapping number of genes between M1 and monocyte subsets in parentheses. The overlap for significance we analysed using the Chi-square test. d) Over Representation Analysis of module M1 using gene sets from the Reactome Pathway database. e) Interaction plot for M1, with gene nodes highlighted.

For LAIV, we identified six distinct co-expression modules (Supp html file 2), which were strongly enriched in genes related to “Diseases associated with O-glycosylation of proteins” (module M1), “Immunoregulatory interactions between a lymphoid and a non-Lymphoid cell” (module M3), “chemokine receptors bind chemokines” (module M4), as well as “interferon signalling” (module M5, Fig S8). Indeed, the hubs of module M5 are well known type I interferon-related genes, such as *ISG15*, *OAS1*, *OASL*, *IFIT1-3*, and *IFITM1*. Altogether, our findings reveal that a strong local antiviral response is elicited in response to LAIV infection.

## Discussion

This study addresses fundamental questions about the immune responses that control and clear Spn carriage and how influenza infection can alter this control. By using for the first time a double experimental human challenge model with LAIV and Spn, we revealed that Spn carriage led to a quick degranulation of pre-existent nasal neutrophils in the human nose and recruitment of monocytes, promoting bacterial clearance. LAIV infection impaired these immune responses following carriage. LAIV is an attenuated influenza strain and wild-type influenza viruses might have even more pronounced effects on the host response to pneumococcus. Carriage in the absence of LAIV was associated with only limited inflammation, corroborating the view of Spn as a commensal bacterium that can asymptomatically colonize healthy adults ^43^. In contrast, robust pro-inflammatory cytokine responses were measured following LAIV at both the protein and gene expression level. Altogether, these results provide explanation for our previous report that LAIV increased acquisition of Spn and carriage density ^14^.

In addition, our findings that LAIV led to impaired blood neutrophil killing capacity and that the addition of TNF-α, which was increased following LAIV, to neutrophils *in vitro* impaired their activity, highlights their crucial roles in susceptibility to secondary bacterial infection ^44^. The association of TIGIT in this impaired neutrophil function following influenza infection warrants further investigation as TIGIT-blocking therapeutics are currently being developed for cancer and HIV treatment ^45^.

We identified IP-10 as a marker for increased susceptibility to Spn and propose this should be further investigated as a potential therapeutic target for secondary bacterial infections associated with virus infections. Our data showed that individuals with higher levels of IP-10 prior to Spn inoculation had higher bacterial density. In a recent study, children with pneumonia with viral and bacterial (predominantly pneumococcal) co-infection had increased levels of IP-10 compared to children with just viral or bacterial pneumonia, which associated with disease severity ^46^. Murine data suggests that IP-10 plays a direct role during pneumonia. Mice with genetic ablation of CXCR3, the receptor for IP-10, CXCL9 and CXCL11, showed increased survival, decreased lung inflammation and less invasion following infection, depending on pneumococcal inoculation strain used ^47^. Moreover, addition of exogenous IP-10 prior to infection of mice with influenza or RSV increased pneumonia severity ^48^.

Our results support previous findings from murine models showing that MCP-1 signalling and monocyte recruitment are key mediators of pneumococcal carriage clearance ^16^. However, contrary to key mechanisms described in murine models, we did not observe any production of IL-17A or neutrophil recruitment to the nose following carriage or associated with carriage clearance ^15–17^, underlining the importance of confirmation of murine findings by human data.

One limitation of this study is that only one pneumococcal serotype 6B isolate was used, future studies using other isolates with a more or less invasive phenotype will be able to answer how generalizable these findings across pneumococcal isolates are. Nonetheless, the observation that carriage density and duration declines in parallel for all serotypes following repeated exposure, suggests that immunological control of newly-acquired Spn is mediated by similar mechanisms independent of the colonizing serotype ^49^.

In conclusion, this study highlighted the importance for innate immunity in the control of carriage density and clearance of Spn, which was impaired by pre-existing viral infections (Fig S9). Secondary bacterial infection following viral respiratory tract infection has a large burden of disease worldwide and disrupting viral-bacterial synergy through host-directed therapy could prove an attractive addition to current therapeutic and vaccination options ^50^.

## Methods

### Study design and sample collection

Healthy adult volunteers were 1:1 randomized to receive either intranasally LAIV (2015/2016 Fluenz Tetra or FluMist Tetra, AstraZeneca, UK) or intramuscular Quadrivalent Inactivated Influenza Vaccination (Fluarix Tetra, GlaxoSmithKline, UK) as described previously ^14^. The control group also received a nasal saline spray, while the LAIV group also received a intramuscular saline injection. Three days post vaccination all subjects were inoculated with 80,000 CFU per nostril of 6B type Spn as described ^6,51^. Nasal microbiopsies (ASL Rhino-Pro©, Arlington Scientific) and nasal lining fluid (Nasosorption™, Hunt Developments) samples were collected and stored at − 80C as previously described ^52^.

### Clinical Trial details

The double-blinded randomized clinical LAIV-EHPC trial was registered on EudraCT (Number 2014-004634-26) on 28^th^ April 2015 and ISRCTN (Number 16993271) on 2^nd^ Sep 2015 and was co-sponsored by the Royal Liverpool University Hospital and the Liverpool School of Tropical Medicine. Key eligibility criteria included: capacity to give informed consent, no immunocompromised state or contact with susceptible individuals, no influenza vaccine or infection in the last two years and not having taken part in EHPC studies in the past three years. The primary endpoint was the occurrence of pneumococcal colonisation determined by the presence of pneumococcus in nasal wash samples (NW) at any time point post inoculation up to and including day 29, detected using classical microbiology or *lytA* qPCR as described ^6,51,53^. In this study, 130 volunteers were inoculated with pneumococcus, giving an 80% power to identify a 50% increase in carriage acquisition. Of 130 vaccinated volunteers, five were natural pneumococcal carriers (two in LAIV arm and three in control arm) and were excluded from further analysis. Another 8 subjects in the LAIV arm were excluded following a systematic LAIV dispensing error by a single practitioner, as recommended by the trial steering group. This resulted in a final 55 subjects analysed in the LAIV arm and 62 subjects in the control arm. Key secondary endpoints included the density of pneumococcal colonisation in NW at each time point following pneumococcal inoculation (days 2, 7, 9, 14, 22 and 29), detected using classical microbiology, the area under the curve of pneumococcal colonisation density following pneumococcal inoculation (days 2, 7, 9, 14, 22 and 29), detected using classical microbiology or by molecular methods (*lytA*), and the immunological mechanisms associated with altered susceptibility to pneumococcus following LAIV. The outcomes reported in this manuscript were a priori included in the study protocol.

### Ethics statement

All volunteers gave written informed consent and research was conducted in compliance with all relevant ethical regulations. Ethical approval was given by the East Liverpool NHS Research and Ethics Committee (REC)/Liverpool School of Tropical Medicine (LSTM) REC, reference numbers: 15/NW/0146 and 14/NW/1460 and Human Tissue Authority licensing number 12548.

### Flow cytometry analysis

Immunophenotyping of nasal cells obtained by curettes was performed as described ^52^. In brief, cells were dislodged from curettes and stained with LIVE/DEAD^®^ Fixable Aqua Dead Cell Stain (ThermoFisher) and an antibody cocktail containing Epcam-PE, HLADR-PECy7, CD16-APC, CD66b-FITC (all Biolegend), CD3-APCCy7, CD14-PercpCy5.5 (BD Biosciences) and CD45-PACOrange (ThermoFisher). Whole blood was stained for 15 min at room temperature with TIGIT-PECy7 (Biolegend) and CD16-APC, followed by 2× 10 min incubation steps with FACSLysis buffer (BD Biosciences) to remove erythrocytes. Samples were acquired on a LSRII flow cytometer (BD) and analysed using Flowjo X (Treestar). Fluorescent minus one controls for each of the included antibodies were used to validate results. For the LAIV and control cohorts, but not the additional validation cohort (Fig S5C), 84/553 samples (15.2%) with less than 500 immune cells or 250 epithelial cells were excluded from further analysis.

### Neutrophil opsonophagocytic killing

Neutrophil killing capacity was evaluated as previously described with minor modifications ^54^. Briefly, neutrophils were isolated through density gradient centrifugation, followed by 45 min incubation with serotype 6B pneumococci (inoculation strain, MOI 100:1), baby rabbit complement (Mast Group, Bootle, UK) and human intravenous immunoglobulin (IVIG; Gamunex, Grifols Inc, Spain). In some experiments, recombinant TNF-α or IP-10 (Biotechne) was added.

### Luminex analysis of nasal lining fluid or stimulated nasal cells

Nasal cells collected in RPMI containing 1% penicillin/streptomycin/neomycin (ThermoFisher) and 10% heat-inactivated FBS (ThermoFisher) were incubated with 50ug/mL DNAse I (Sigma Aldrich) at room temperature for 20 min and filtered over a 70um filter (ThermoFisher). Cells were spun down at 440xg for 5 min, resuspended, counted and incubated at 250,000 cells/mL in 96-wells or 384-wells plates (ThermoFisher). Heat-killed Spn inoculation strain was added at a concentration of 5 ng/mL of total protein (corresponding to 4.3×10^7 CFU/mL) and cells were incubated for 18 h. Bacterial protein concentration was measured by Bradford assay, using bovine serum albumin as standard and titration experiments were performed to determine dose. Supernatant was collected and stored at −80C until analysis. For nasosorption filters, cytokines were eluted from stored filters using 100 uL of assay buffer (ThermoFisher) by centrifugation, then the eluate was cleared by further centrifugation at 16,000g. Prior to analysis, samples were centrifuged for 10 min at 16,000xg to clear samples. These were acquired on a LX200 using a 30-plex magnetic human Luminex cytokine kit (ThermoFisher) and analysed with xPonent3.1 software following manufacturer’s instructions. Samples were analysed in duplicates and nasosorption samples with a CV > 25% were excluded.

### RNA extraction and sequencing

Nasal cells were collected in RNALater (ThermoFisher) at −80C until extraction. Extraction was performed using the RNEasy micro kit (Qiagen) with on column DNA digestion. Extracted RNA was quantified using a Qubit™ (ThermoFisher). Sample integrity assessment (Bioanalyzer, Agilent), library preparation and RNA-sequencing (Illumina Hiseq4000, 20M reads, 100 paired-end reads) were performed at the Beijing Genome Institute (China).

### Nanostring

Purified blood neutrophils were stored in RLT buffer (Qiagen) with 1% 2-mercaptoethanol (Sigma) at −80C until RNA extraction as above. The single cell immunology v2 kit (Nanostring) was used with 20 pre-amp cycles for all samples. Hybridized samples were prepared on a Prep Station and scanned on a nCounter® MAX (Nanostring). Raw counts were analysed using DESeq2 using internal normalization, which gave lower variance than normalizing to included housekeeping genes. DEG were identified using a model matrix correcting for repeated individual measurements. Log counts per million (CPM) from raw counts were calculated using the ‘edgeR’ package.

### RNA sequencing analysis

Quality control of raw sequencing data was done using fastQC. Mapping to a human reference genome assembly (GRCh38) was done using STAR 2.5.0a ^55^. Read counts from the resulting BAM alignment files were obtained with featureCounts using a GTF gene annotation from the Ensembl database ^56,57^. The R/Bioconductor package DESeq2 was used to identify differentially expressed genes among the samples, after removing absent features (zero counts in more than 75% of samples) ^58^. Genes with an FDR value < 0.1 and an absolute fold change (FC) > 1.5 were identified as differentially expressed.

### Co-expression analysis

For co-expression analysis, counts were normalized using log CPM and the log2 fold change was calculated for each time point in a subject-wise manner. The co-expression analysis was performed separately for each group (control and LAIV) using the CEMiTool package developed by our group and available at Bioconductor (https://bioconductor.ora/packaaes/release/bioc/html/CEMiTool.html) ^39^. This package unifies the discovery and the analysis of coexpression gene modules, evaluating if modules contain genes that are over-represented by specific pathways or that are altered in a specific sample group. A p-value = 0.05 was applied for filtering lowly expressed genes

### Statistical analysis

All experiments were performed randomised and blinded. Two-tailed statistical tests are used throughout the study. When log-normalized data was not normally distributed, non-parametric tests were performed and multiple correction testing (Benjamin-Hochberg) was applied for gene expression and Luminex analysis.

### Data availability

Raw RNA-sequencing data have been deposited in the GEO repository. All other underlying data is provided in the manuscript

## Acknowledgements

DF is supported by the Medical Research Council (grant MR/M011569/1), Bill and Melinda Gates Foundation (grant OPP1117728) and the National Institute for Health Research (NIHR) Local Comprehensive Research Network. Flow cytometric acquisition was performed on a BD LSR II funded by a Wellcome Trust Multi-User Equipment Grant (104936/Z/14/Z). SJ received support from the Royal Society of Tropical Medicine and Hygiene for Nanostring analysis. HIN is supported by FAPESP (grants 2013/08216-2 and 2012/19278-6). We would like to thank all volunteers for participating in this study and Rachel Robinson, Cath Lowe, Lepa Lazarova, Katherine Piddock and India Wheeler for clinical support. We would also like to acknowledge Stephen Gordon and Michael Mina for their input.

## Author contributions

SPJ contributed to conceiving, designing, performing and analysing experiments and writing of the paper. FM and HIN contributed analysing experiments and writing of the paper. BFC, MH, EM, ES, JFG, CS, JR, SP, EN, ELG, WAASP, DB contributed to conducting and analysing experiments. AHW, HH, CH, HA, SZ, VC, JR contributed to sample collection and design of the study. DMF contributed to conceiving, designing and analysing experiments, design of the study and writing of the paper. All authors have read and approved the manuscript.

---

**Supplementary Figure 1.**
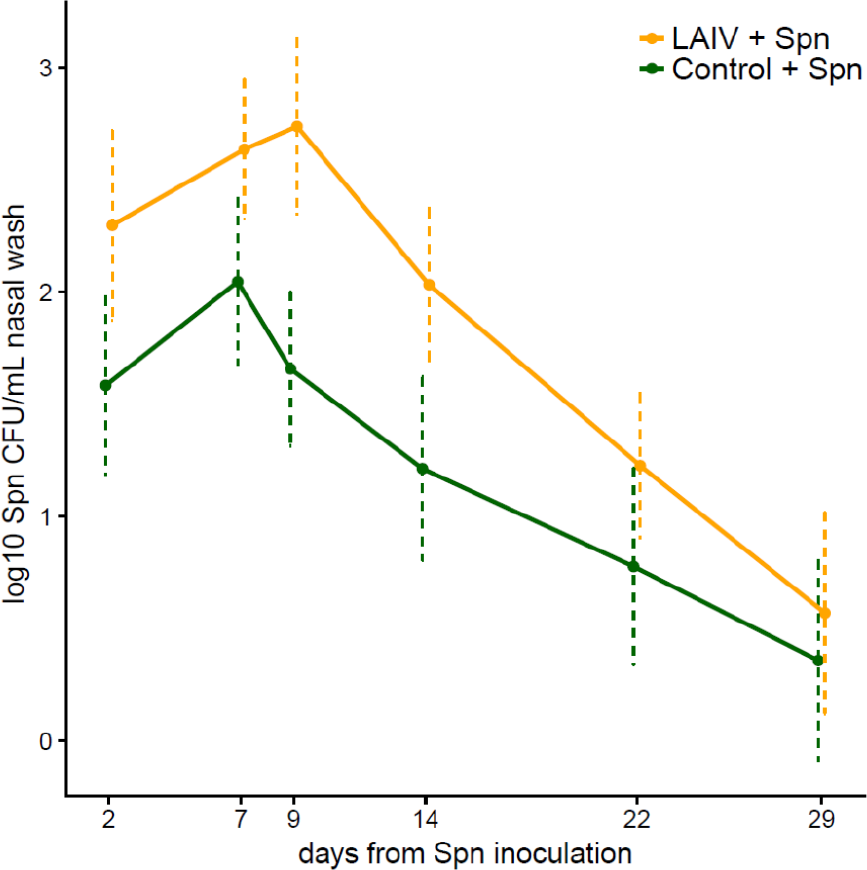
Pneumococcal density in the LAIV and Control cohorts. Mean and standard error of log-transformed carriage density [CFU/mL nasal wash] for carriage-positive volunteers (defined by detection of Spn at any timepoint) are shown for the Spn group (green, n = 14-24 samples per time-point) and LAIV + Spn group (orange, n = 16-25). Density for samples with undetectable pneumococcal load was set at 0.1 CFU/mL nasal wash.

**Table S1.**
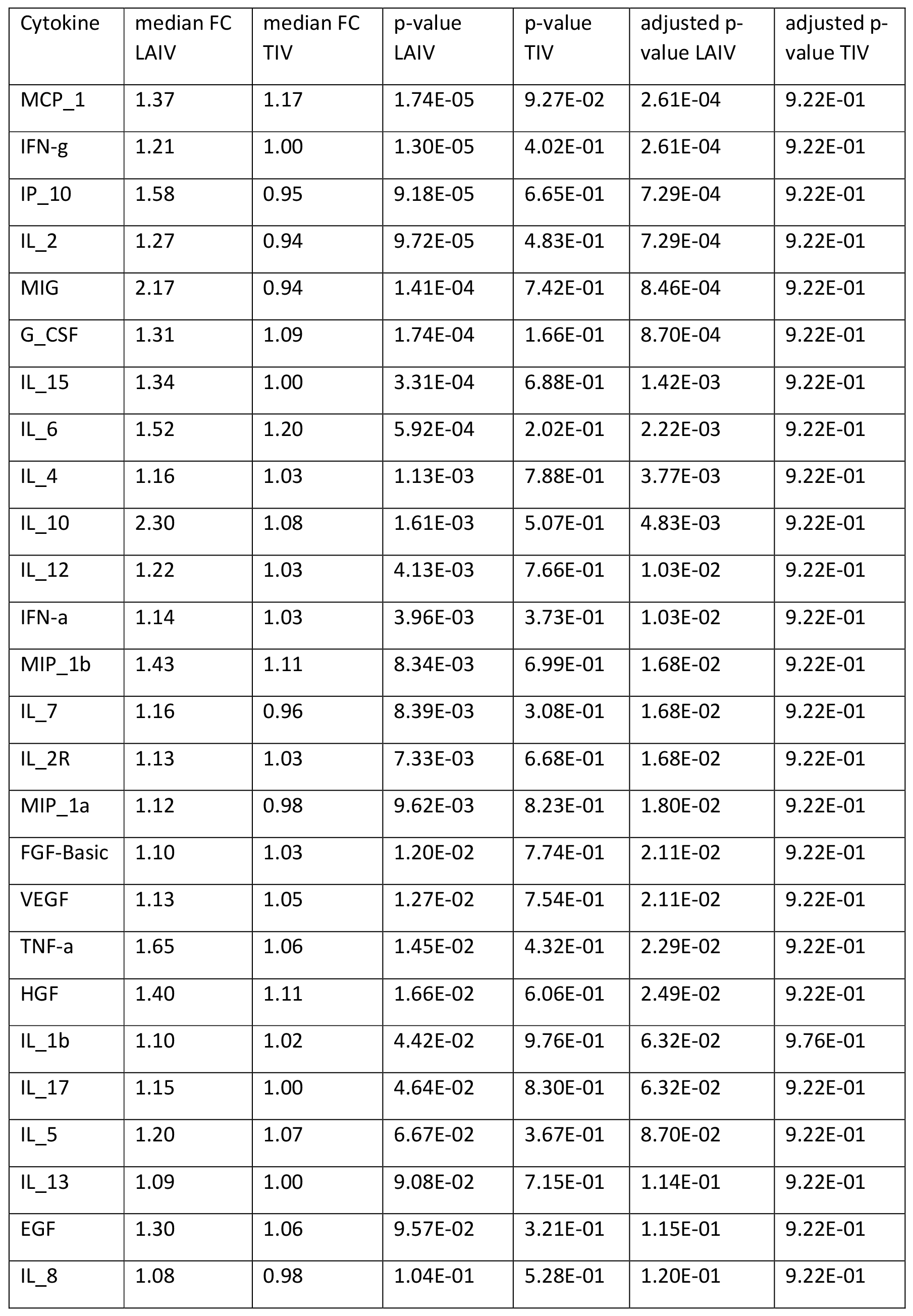
Cytokine induction following LAIV or control vaccination, prior to Spn inoculation. Levels of 30 cytokines in nasal fluid were measured before and 3 days after vaccination, just prior to pneumococcal challenge. Median FC, Wilcoxon p-values, and FDR-corrected p-values are depicted. N = 38 per group.

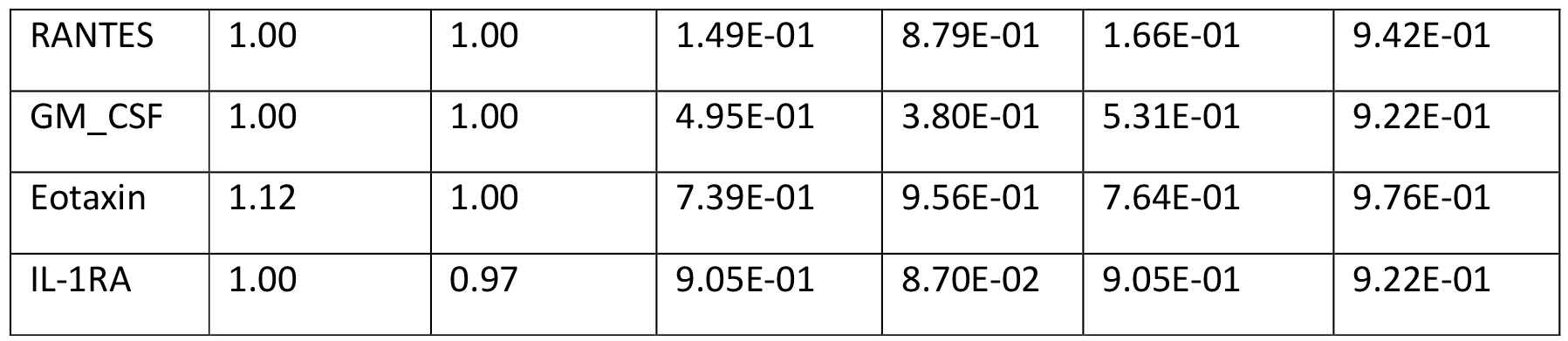

**Supplementary Figure 2.**
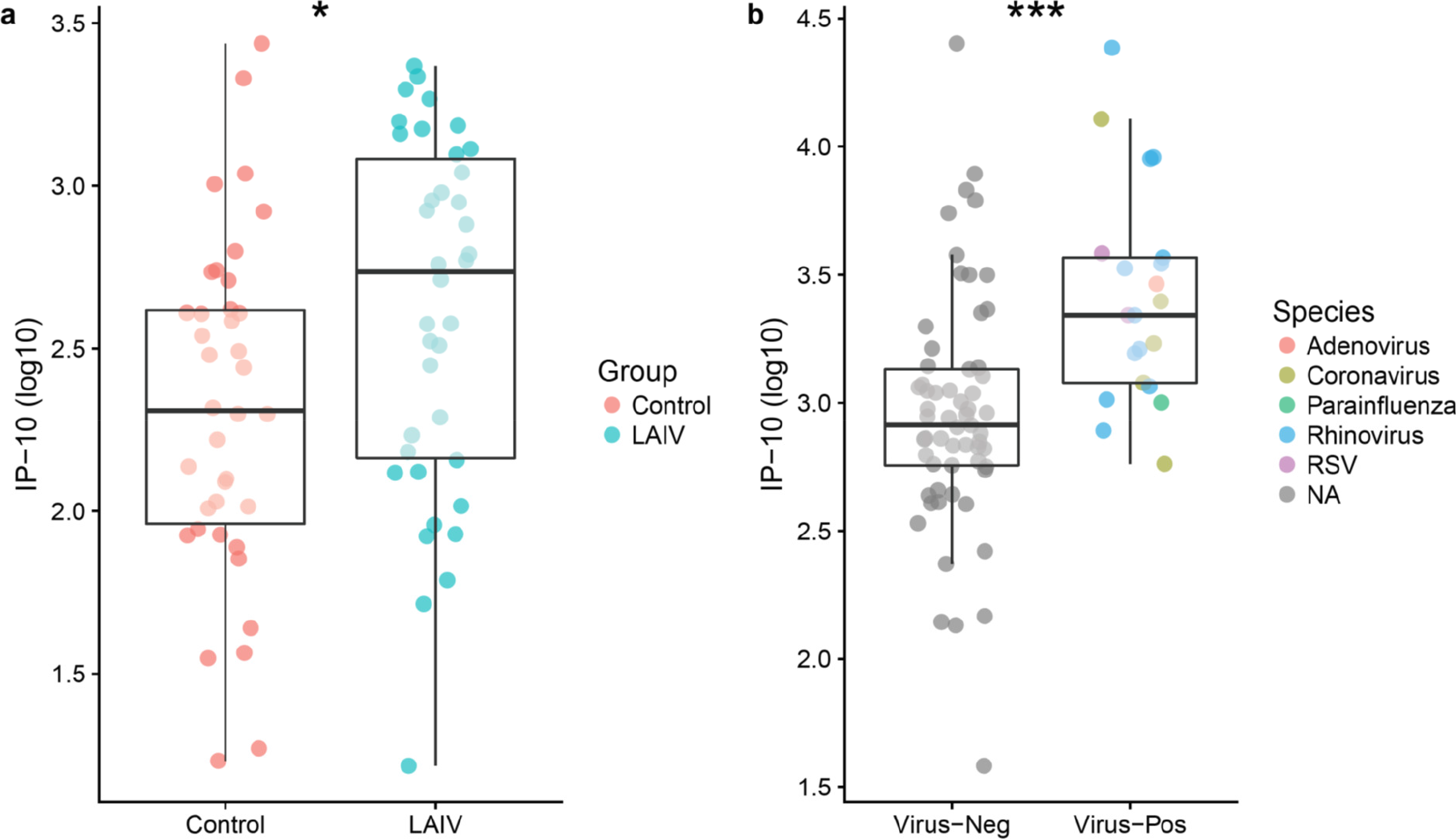
Levels of IP-10 are increased by virus infection. a) Levels of IP-10 at day 0 in the LAIV cohort, measured by Luminex in nasosorption (n=77). b) Levels of IP-10 at day-5 to Spn inoculation in a cohort with known viral URT state measured by ELISA (n=82). Oropharyngeal swabs collected 5 days before Spn inoculation were assessed for viral multiplex PCR panel for detection of influenza A and B (n=0), respiratory syncytial virus (n=2), human metapneumovirus (n=0), human rhinovirus (n=12), parainfluenza viruses 1–4 (n=1), and coronaviruses OC43, NL63, 229E, and HKU1 (n=5). Individual volunteers are shown and median and interquartile ranges are depicted per group. * p < 0.05, *** p < 0.001 by Mann-Whitney test.

**Supplementary Figure 3.**
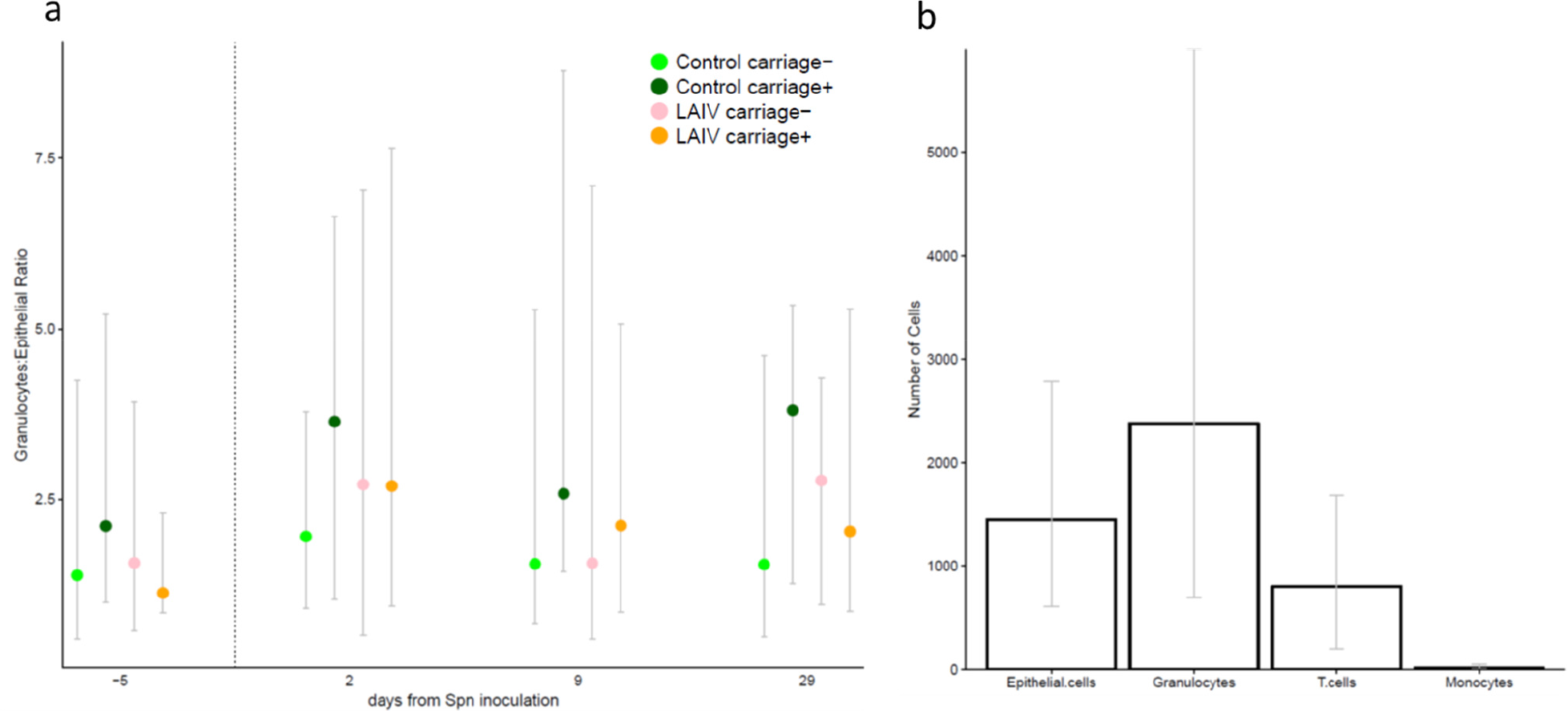
Levels of a) granulocytes, predominantly neutrophils based on CD16 expression, at the nasal mucosa. Median and interquartile range are shown for cell numbers normalized to numbers of epithelial cells. The x-axis shows days relative to inoculation. b) Absolute numbers of cells observed per nasal sample (median and IQR are shown for 117 subjects at baseline).

**Table S2.** List of differentially expressed genes (DEG with p < 0.01) in sorted blood neutrophils 3 days following LAIV vaccination compared to pre-vaccination and controls, correcting for repeated individual measurements.

| Gene | log2<br>FC | p-value | padj |
| --- | --- | --- | --- |
| MAP4K2 | 1.67 | 3.23E-05 | 1.18E-02 |
| HLA-DPB1 | -3.28 | 5.87E-04 | 1.07E-01 |
| IL1R2 | -1.35 | 2.13E-03 | 2.34E-01 |
| S100A9 | -2.23 | 2.57E-03 | 2.34E-01 |
| CASP2 | -1.31 | 3.67E-03 | 2.67E-01 |
| PSMC2 | -1.55 | 4.70E-03 | 2.85E-01 |
| TFRC | 2.18 | 7.26E-03 | 3.09E-01 |
| POLR2A | -1.24 | 8.29E-03 | 3.09E-01 |
| TIGIT | 1.84 | 8.30E-03 | 3.09E-01 |
| ITGAE | -1.54 | 8.48E-03 | 3.09E-01 |

**Supplementary Figure 4.**
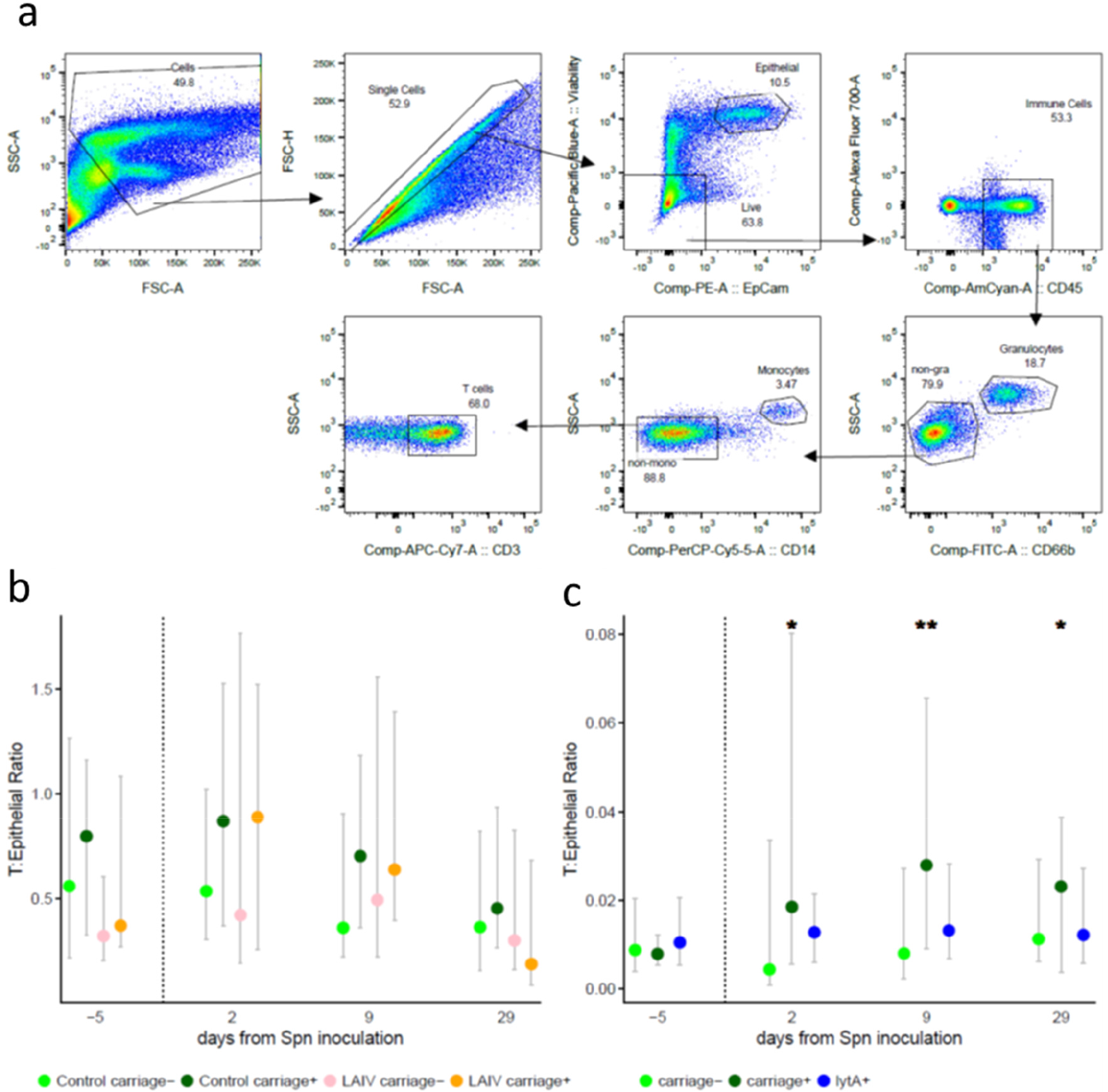
a) Gating strategy of nasal cells for one representative volunteer. b) Levels of T cells at the nasal mucosa. Median and interquartile range are shown for cell numbers normalized to numbers of epithelial cells. The x-axis shows days relative to inoculation. c) Median and interquartile range of monocytes levels in the control group over time with carriage detected by classical microbiology (dark green, carriage+), molecular methods only (blue, *lytA+*) or not at all (light green, carriage-). *p < 0.05, **p < 0.01 by Wilcoxon paired test.

**Supplementary Figure 5.**
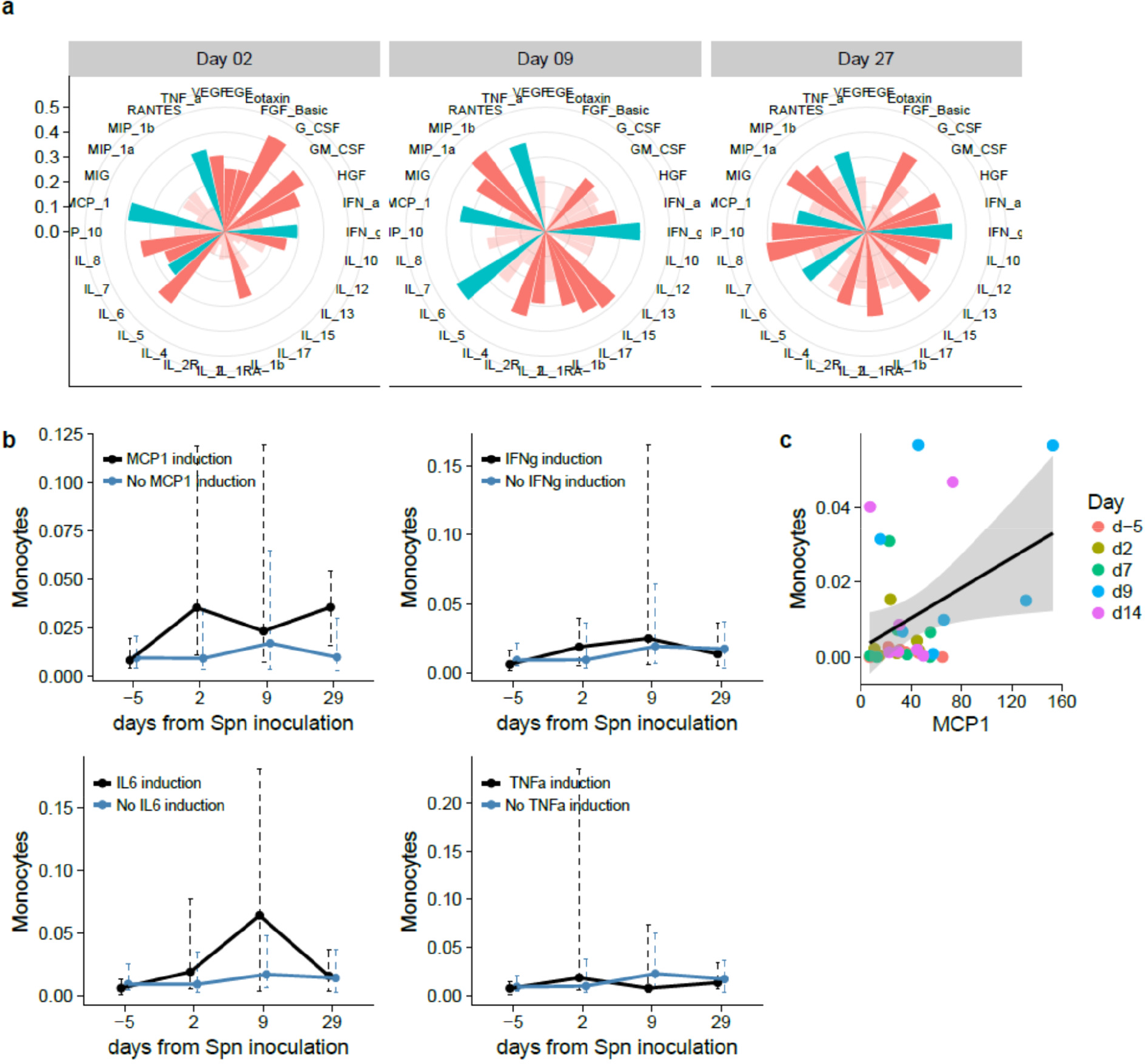
Monocyte recruitment associates with MCP1 levels and is reproducible in an independent volunteer cohort. a) Spearman’s correlation between the 30 measured cytokines and nasal monocytes levels for each timepoint. The length of the bar depicts the rho value, cyan bars represent cytokines that correlate significantly at all days, red bars associate for a specific day, and transparent red bars are not significantly associated for that day. b) Monocyte levels over time in those volunteers who most upregulate MCP1, IFNg, IL6 or TNFa at day 2 (top quartile FC induction versus below top quartile) or not. * p < 0.05 by MW-test of AUC of inducers and non-inducers). c) Levels of monocytes and MCP1 in a second, independent cohort of 7 carriers without vaccination. Symbols represent individual subjects, with color corresponding to day relative to Spn inoculation. Effect of MCP1 on monocyte levels by generalized linear regression analysis are shown, correcting for repeated individual measurements.

**Table S3.**
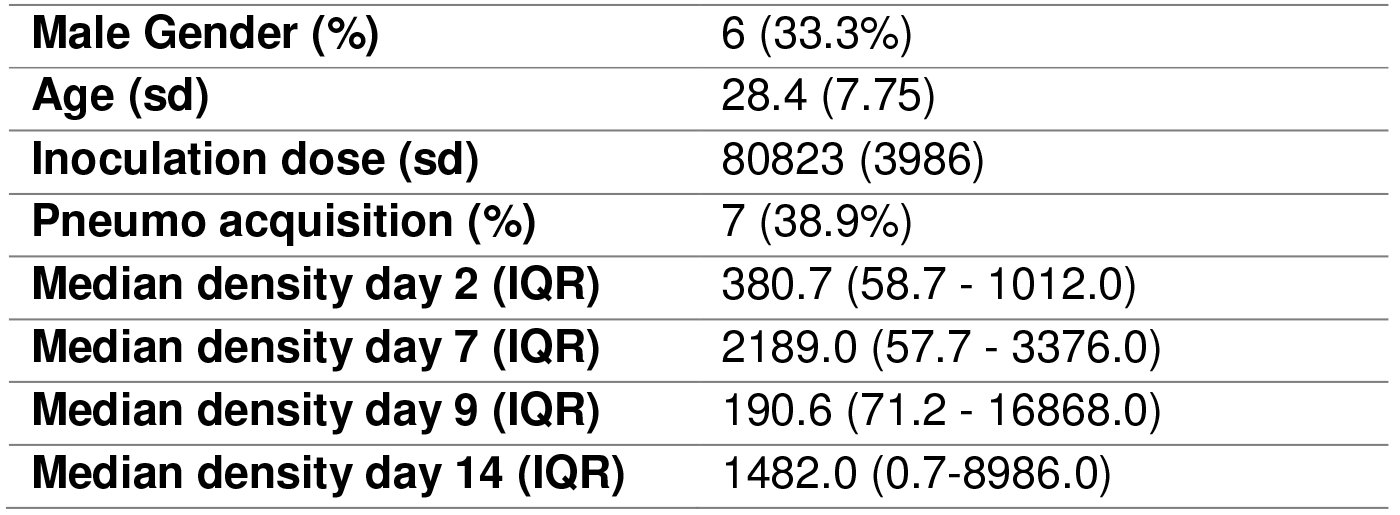
Participant characterization of the validation cohort (n=18).

|  |  |
| --- | --- |
| **Male Gender (%)** | 6 (33.3%) |
| **Age (sd)** | 28.4 (7.75) |
| **Inoculation dose (sd)** | 80823 (3986) |
| **Pneumo acquisition (%)** | 7 (38.9%) |
| **Median density day 2 (IQR)** | 380.7 (58.7 - 1012.0) |
| **Median density day 7 (IQR)** | 2189.0 (57.7 - 3376.0) |
| **Median density day 9 (IQR)** | 190.6 (71.2 - 16868.0) |
| **Median density day 14 (IQR)** | 1482.0 (0.7-8986.0) |

**Supplementary Figure 6.**
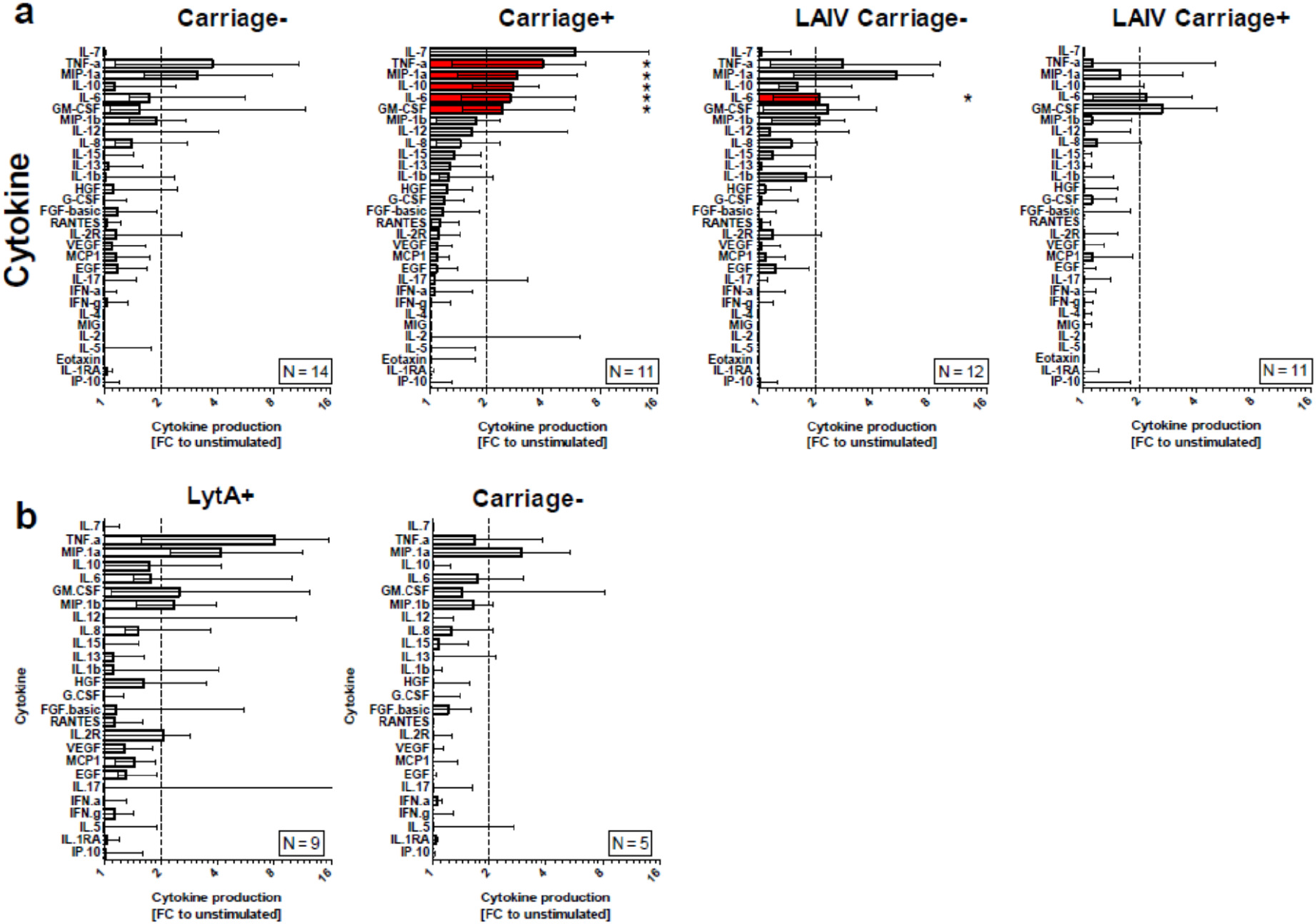
Whole nasal cells were collected 29 days post Spn inoculation and stimulated for 18 hours with heat-killed pneumococcus. Supernatant was collected and levels of 30 cytokines were measured by Luminex. Four cytokines were not measured above the limit of detection (IL-2, IL-4, MIG and Eotaxin) and these were excluded from analysis. a) Responses for all measured cytokines for the 4 groups are shown. b) Production in carriage-volunteers subdivided in those who had very low density carriage detectable only by molecular methods (*lytA+*) or not at all (carriage-). The dotted line indicates a 2-fold increase over the unstimulated control. Median and interquartile range of FC to unstimulated are shown *, Significantly induced cytokines are shown in red. (FC > 2, q < 0.05, Wilcoxon followed by Benjamini-Hochberg (BH) correction).

**Supplementary Figure 7.**
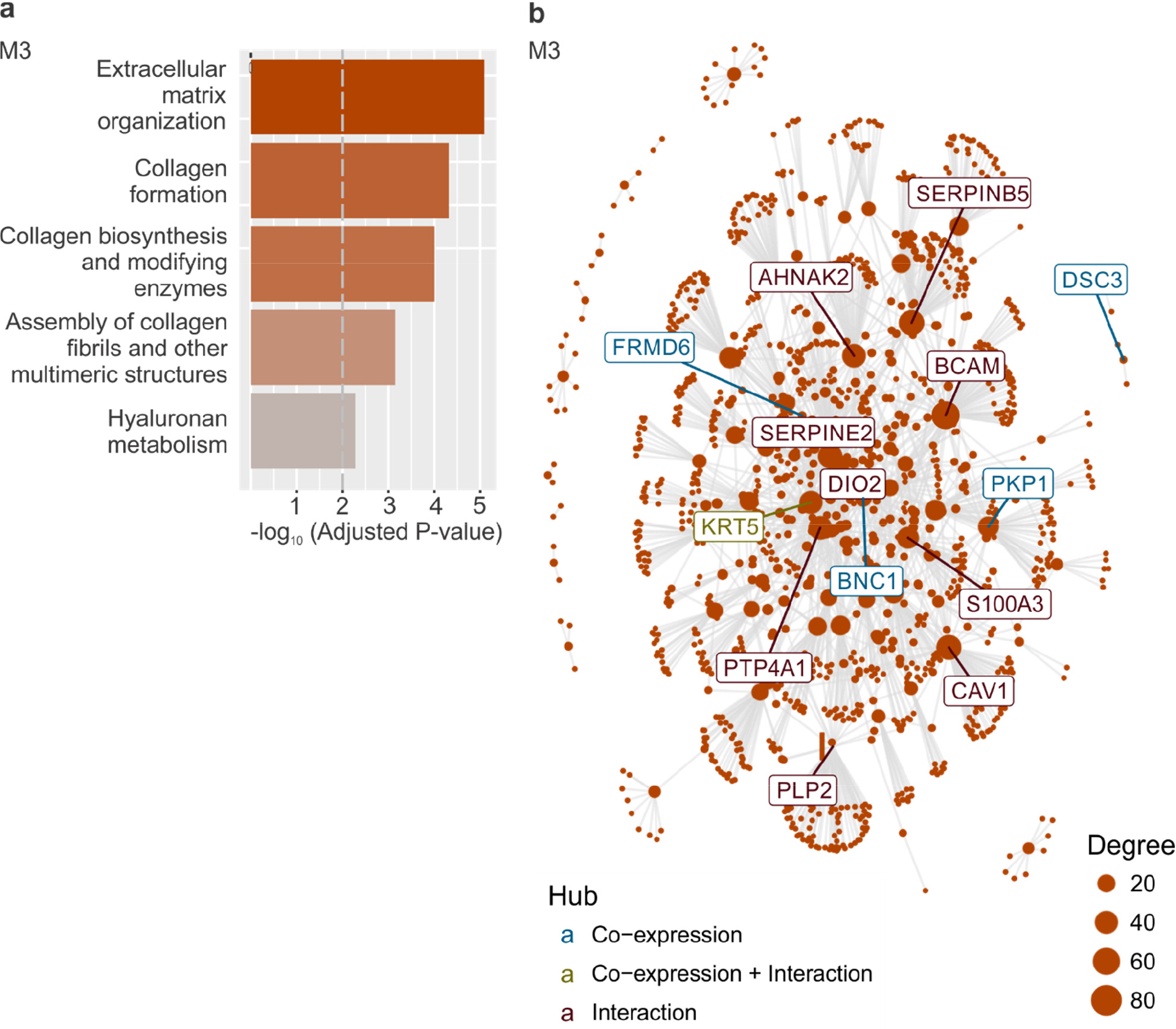
CEMiTool applied to control group – M3. Raw counts were normalized using logCPM and log2-fold change were calculated for each timepoint against the baseline after which coexpression modules were extracted. a) Over Representation Analysis of module M3 of the control group using gene sets from the Reactome Pathway database. e) Interaction plot for M3, with gene nodes highlighted.

**Supplementary Figure 8.**
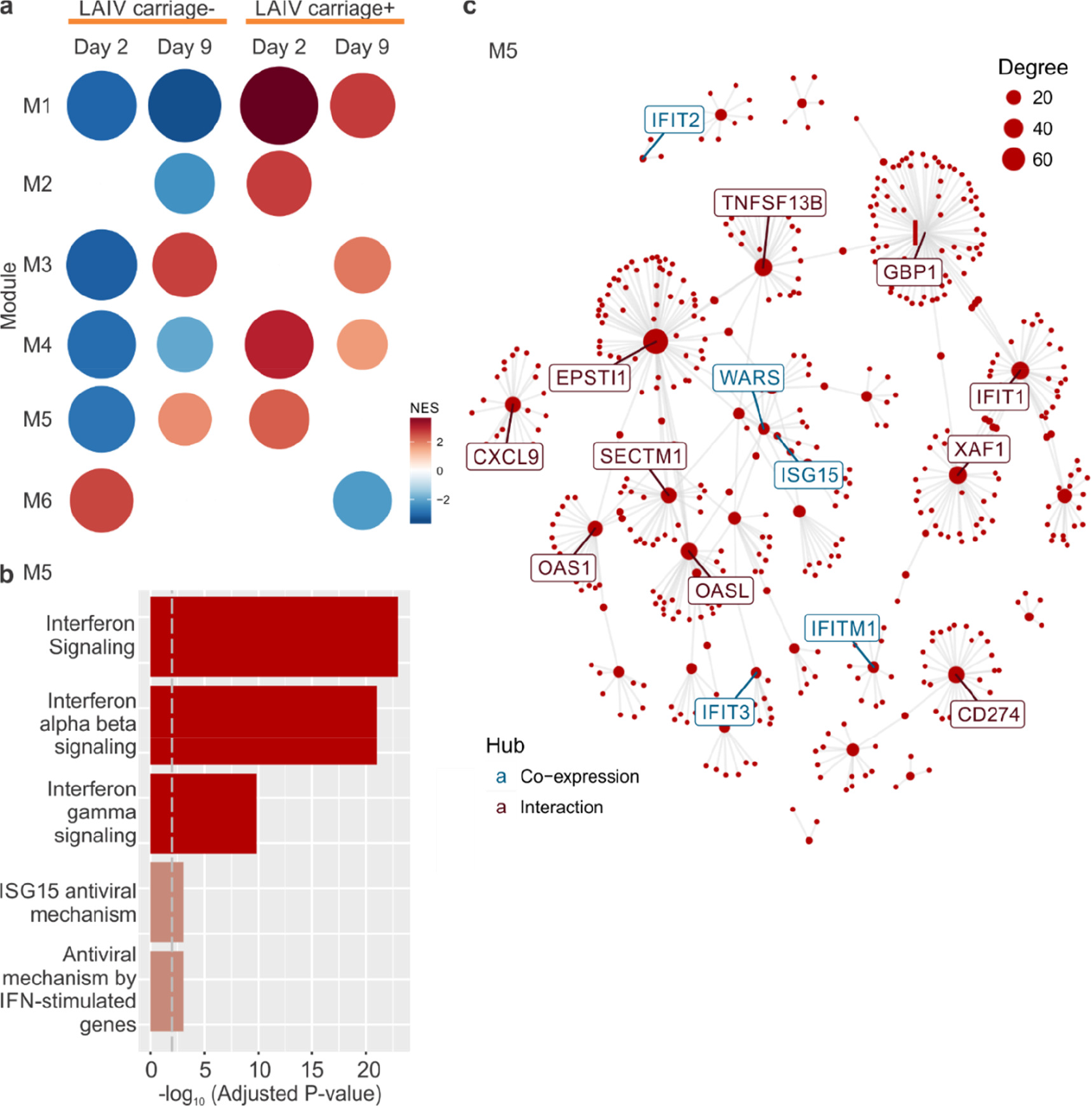
CEMiTool applied to LAIV. Raw counts were normalized using logCPM and log2fold change were calculated for each timepoint against the baseline after which co-expression modules were extracted. a) Gene Set Enrichment Analyses showing the module activity on each timepoint for carriage+ and carriage-LAIV groups. b) Over Representation Analysis of module M5 of the LAIV group using gene sets from the Reactome Pathway database. e) Interaction plot for M5, with gene nodes highlighted.

**Supplementary Figure 9.**
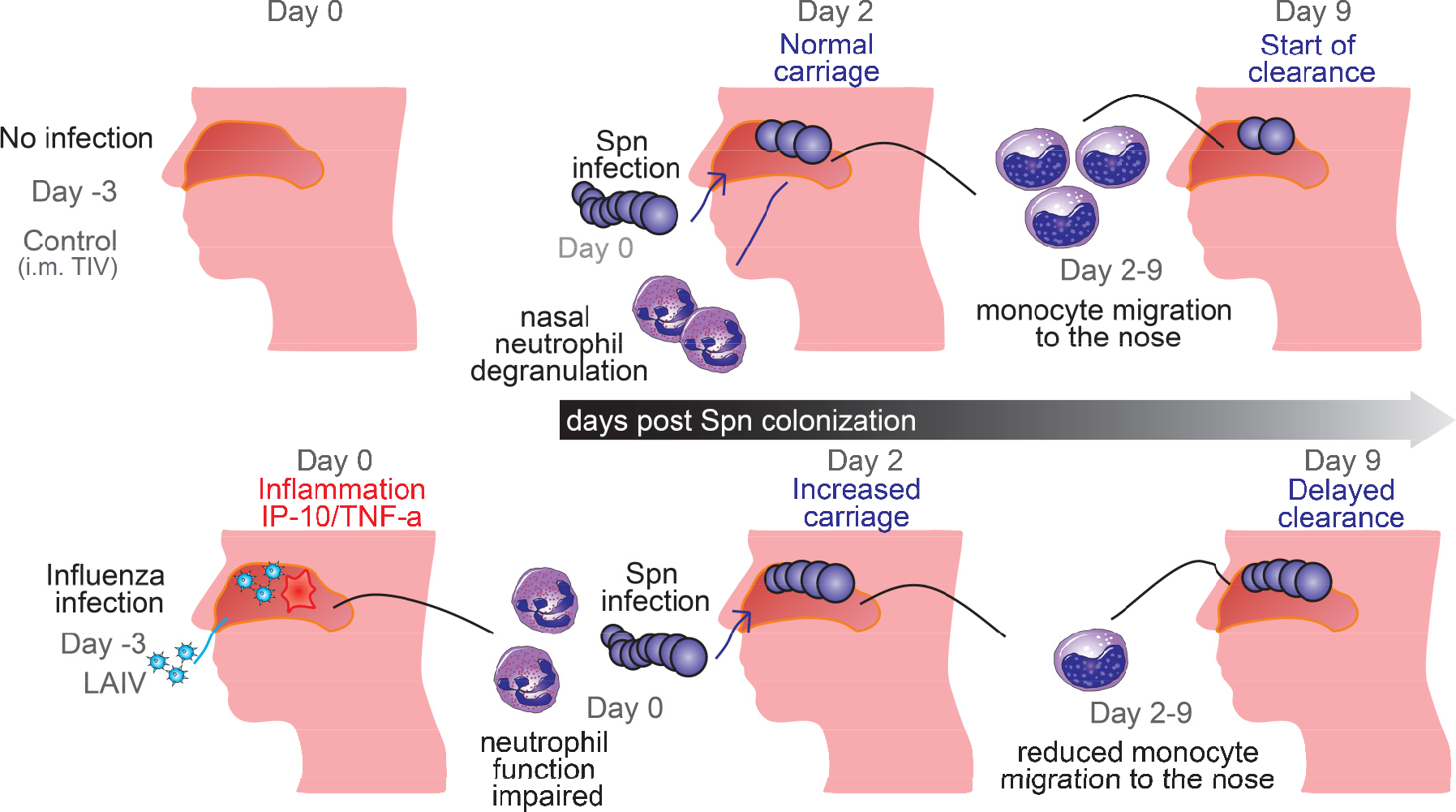
Summary of immune mechanisms associating with control of Spn carriage and their disruption by LAIV. Carriage in the absence of influenza leads to quick degranulation of nasal neutrophils followed by a recruitment of monocytes to the nose, associating with the start of clearance. Influenza infection leads to inflammation, impairing this innate control of carriage.

